# Niche dimensionality drives microbial community structure

**DOI:** 10.1101/2025.11.02.686095

**Authors:** Karthik Srinivasan, German Plata, Purushottam D. Dixit

## Abstract

Niche dimensionality links environmental complexity to ecosystem structure. Although niche theory is often invoked in investigations of microbiomes, most models assume very high-dimensional coexistence, effectively sidestepping the role of dimensionality. However, direct estimates of niche dimensionality of microbiome have been lacking. Here we use joint species distribution modeling (JSDM) to infer niche dimensionality from relative abundance data alone. In paired 16S rRNA–metabolomics datasets, inferred dimensionality closely tracked the complexity of the metabolic environment, validating our abundance-only approach. Across nearly 200 human gut microbiome studies, lower dimensionality coincided with greater inter-species competition (metabolic niche overlap) and reduced biodiversity, consistent with the macroecological *niche dimensionality hypothesis*. Consumer/resource simulations reproduced these empirical relationships when both species-intrinsic metabolic tradeoffs and species-extrinsic environmental tradeoffs were imposed, with the latter dominating. Dimensionality was reduced in stressed or diet-simplified microbiomes and correlated broadly with ecological stress markers; negatively with prevalence of generalists and positively with microbial load. Together, these results establish niche dimensionality as a measurable and previously overlooked driver of microbial community structure.

## I. INTRODUCTION

Host-associated microbiomes are extraordinarily complex ecosystems comprising thousands of microbial taxa spanning multiple kingdoms of life^1,2^. These communities are essential to human health^3^ and animal productivity^4^, yet the ecological principles governing their assembly, stability, and function remain incompletely understood.

Niche theory provides a quantitative framework linking environmental variables to community structure across scales^5,6^. A species’ niche is the multidimensional set of environmental conditions that supports its persistence^7^; coexistence of multiple species is facilitated when they occupy distinct regions of this space, thereby minimizing *niche overlap* (inter-species competition)^5,6,8^. In microbial systems, niche theory is often invoked to explain observed biodiversity via mechanisms such as niche partitioning^9–13^ and niche construction^14–16^.

The extent of niche overlap is fundamentally constrained by niche dimensionality (*η*_D_ from here onward), the number of independent environmental axes governing community composition^17,18^, with higher niche dimensionality leading to a reduction in niche overlap and facilitation of biodiversity and vice versa for lower dimensional niches^8,18–21^. Empirical studies in macroecology have additionally linked *η*_D_ to community stability, invasibility, and evolutionary diversification^17,22–31^, underscoring its foundational role in ecosystem organization.

Species’ environments are invariably multidimensional and correlated, and species’ responses to environmental variables are heterogeneous and often unknown. Therefore, estimation of niche dimensionality via cataloging all *relevant* niche axes is rarely feasible. Ecologists therefore infer an *effective* niche dimensionality from empirical data on traits, interaction networks, or trophic structure^32–38^. These estimates treat niche dimensionality as an emergent property, the number of effective independent ecological axes required to reproduce observed community patterns.

To date, estimates of niche dimensionality have not been extended to microbial ecosystems, likely because trait information is limited and controlled competition experiments are rarely feasible. Paradoxically, however, most theoretical models of microbiomes implicitly or explicitly assume very high-dimensional niche spaces, often manifested as statistically independent interaction coefficients^39^ in Lotka–Volterra or consumer–resource formulations^40–49^, thereby obscuring the potential role of niche dimensionality constraints in shaping community structure.

To address this gap, we develop a statistical approach to infer microbial niche dimensionality directly from community composition using joint species distribution modeling (JSDMs)^50–53^. JSDMs traditionally relate measured environmental covariates to species abundances^50^. Their latent-variable extensions^53–56^ introduce latent factors that capture unmeasured and unknown environmental influences. Here, we extend the use of JSDMs to estimate effective niche dimensionality *η*_D_ from relative abundances by decomposing taxa into ecosystem-specific latent “resources” and species-specific preferences toward those resources; *η*_D_ is identified as the characteristic dimension of the latent space required to reconstruct observed community composition.

In paired 16S rRNA–metabolomics data^57^, *η*_D_ tracked the dimensionality of the metabolic environment, validating that abundance data encode niche structure. Applied to ∼ 10^5^ samples spanning nearly 200 gut microbiome studies^58^, inferred *η*_D_ was consistently very low (median *η*_D_ = 4), indicating co-existence in surprisingly low-dimensional niche spaces. Across studies, reduced *η*_D_ was associated with higher metabolic niche overlap (a proxy for competition) and lower biodiversity, consistent with the macroecological *niche dimensionality hypothesis*^25–27,59^. Consumer–resource simulations recapitulated these empirical patterns only when both species-intrinsic metabolic tradeoffs^9,11^ and species-extrinsic environmental tradeoffs were imposed, with the latter exerting the dominant influence. Additionally, we found that niche dimensionality was reduced in stressed or diet-simplified microbiomes and broadly covaried with ecological stress markers; negatively with prevalence of generalists^60^ and positively with microbial load^61^. Together, these results identify niche dimensionality as an empirically measurable, previously overlooked driver of microbial ecosystem organization.

## II. RESULTS

We first introduce and validate a latent-variable joint species distribution model (JSDM) to infer *η*_D_ from relative abundance data. We then show that inferred dimensionality mirrors environmental complexity and follows macroecological expectations linking dimensionality, competition, and diversity. Next, we identify the mechanistic origins of low dimensionality using consumer–resource simulations with intrinsic (metabolic) and extrinsic (environmental) tradeoffs. Finally, we demonstrate that host diet and health systematically modulate microbial niche dimensionality.

### A. Latent variable JSDM quantifies niche dimensionality

To estimate microbial niche dimensionality from relative abundances, we employ a latent-variable JSDM^50–53^ formulated as a non-linear low-rank factorization^54–56^ (see Appendix I):

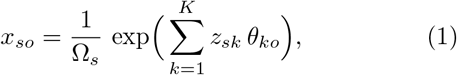

where *x*_*so*_ is the relative abundance of taxon *o* in sample *s* (Ω_*s*_ is the normalization), *z*_*sk*_ are sample-specific latent embeddings (effective “resources”), and *θ*_*ko*_ are speciesspecific loadings (“preferences”). Notably, we have previously shown that this particular form of the JSDM can be derived from the consumer/resource model wherein latent variables *z*_*sk*_ represent time averages of nutrient concentrations^56^.

Traditionally, JSDMs relate measured environmental variables to species abundances^50^. The latent variable formulation instead infers *effective* ecological axes directly from abundance data, capturing both measured and unmeasured influences on community composition^51–56^. The inferred latent variables *z*_*sk*_ need not map one-to-one to individual metabolites or abiotic factors. Instead, they represent effective environmental features that combine external inputs and microbe-driven niche construction.

We assessed ecological interpretability of the JSDM using paired 16S rRNA–metabolomics data^57^. After fitting Eq. 1, we (i) mapped bacterial taxa to ∼ 7000 genomescale gut microbial models^62^ and imputed metabolite-use preferences *r*_*mo*_ via flux balance analysis (FBA)^63,64^ (Appendix II, III), and (ii) regressed *r*_*mo*_ on species loadings *θ*_*ko*_ (using logistic regression) and metabolite concentrations on ecosystem embeddings *z*_*sk*_ (using linear regression). In both cases, models using the inferred embeddings outperformed shuffled controls (Fig. 1c,d), showing that abundance-only latent factors capture biologically meaningful microbe–environment interactions (Appendix II, III).

**FIG. 1.**
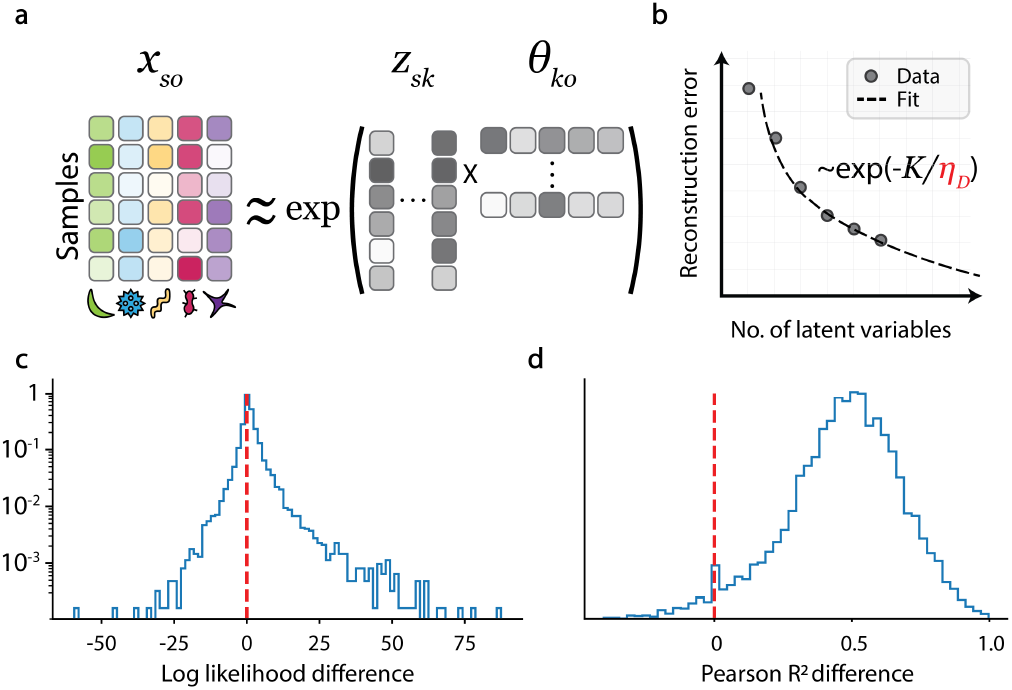
Latent-variable approach to estimate niche dimensionality *η*_D_. **a**, Non-linear low-rank factorization of relative abundances; *z*_*sk*_ are ecosystem-specific latent embeddings (“resources”) and *θ*_*ko*_ are species-specific loadings (“preferences”). **b**, *η*_D_ is the characteristic dimension *K* needed to reconstruct observed compositions with high fidelity. **c**, Species loadings *θ*_*ko*_ predict imputed metabolite-preference profiles (histogram of difference in log-likelihood vs. shuffled). **d**, Ecosystem embeddings *z*_*sk*_ predict measured metabolite concentrations (histogram of difference in Pearson *R*^2^ vs. shuffled).

We define *η*_D_ as the characteristic number of latent dimensions, *K*, required to accurately reconstruct community composition. Specifically, we fit the decay of reconstruction error with increasing *K* using an exponential function and extract its characteristic scale (Appendix I). Our approach parallels prior estimates of niche dimensionality that employed latent spaces to embed ecological structures such as food webs^32,34–37^ or pairwise competition networks^38^. Robustness analyses using alternative dimensionality metrics, namely, the participation ratio and spectral entropy of singular values, produced highly correlated estimates of niche dimensionality (Appendix IV).

### B. Estimated dimensionality reflects environmental complexity and follows the niche dimensionality hypothesis

Metabolism of small molecules plays a dominant role in determining microbial community structure^42,45^. Therefore, we hypothesized that *η*_D_ inferred from taxon abundances alone should mirror the complexity of the metabolic environment. Indeed, in a large collection of paired human gut 16S rRNA–metabolomics datasets^57^, *η*_D_ correlated with the dimensionality of the metabolite concentration space (Spearman *r* = 0.57, *p* = 4.8 × 10^−3^; Fig. 2a), where environmental dimensionality was quantified by singular value decomposition (Appendix II).

**FIG. 2.**
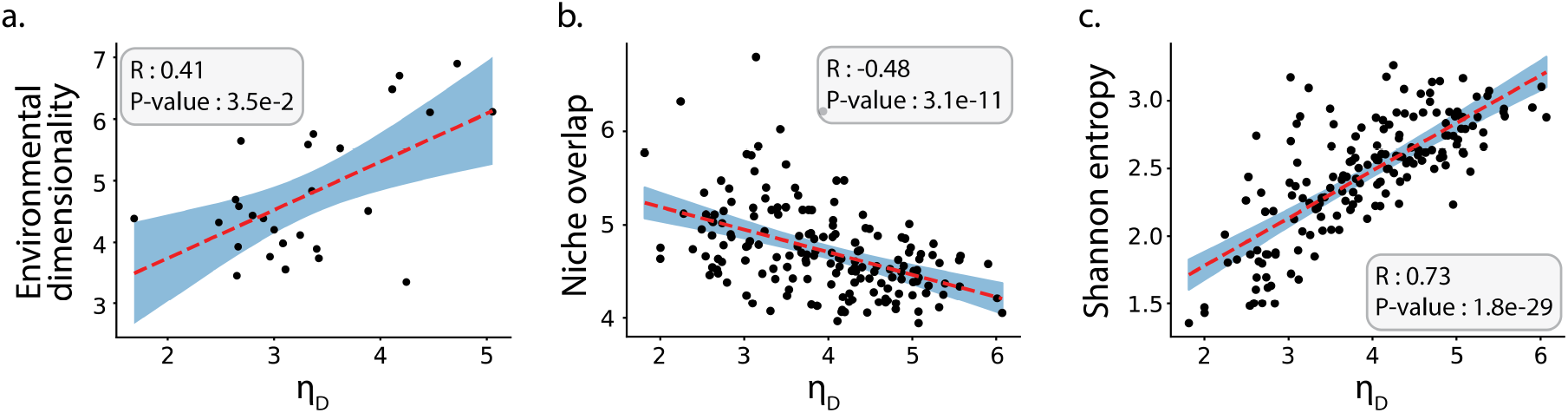
Niche dimensionality captures environmental complexity and macroecological expectations. **a**, *η*_D_ estimated from relative abundances correlates with the dimensionality of the metabolic environment. **b**, *η*_D_ correlates negatively with metabolic niche overlap (competition). **c**, *η*_D_ correlates positively with *α*-diversity (Shannon index).

Since niche dimensionality constrains niche overlap, lower *η*_D_ should coincide with stronger competition and reduced biodiversity^8,17–21^. To investigate these relationships, we estimated *η*_D_ for ∼ 200 gut microbiome studies comprising ∼ 10^5^ samples^58^. Interestingly, *η*_D_ estimates varied significantly across the datasets (median 4.0, coefficient of variation 0.23). Notably, metabolic niche overlap (imputed via FBA; Appendices III, II) correlated negatively with *η*_D_ (Spearman *r* = − 0.48, *p* = 3.1 × 10^−11^; Fig. 2b). Conversely, *η*_D_ correlated positively with *α*-diversity (Spearman *r* = 0.73, *p* = 1.8 × 10^−29^; Fig. 2c). The negative association between *η*_D_ and niche overlap remained statistically significant even after controlling for *α*-diversity (partial Spearman *r* = − 0.26, *p* = 6.8×10^−4^; Fig. 3), confirming that dimensional constraints on competition are not an artifact of community diversity.

**FIG. 3.**
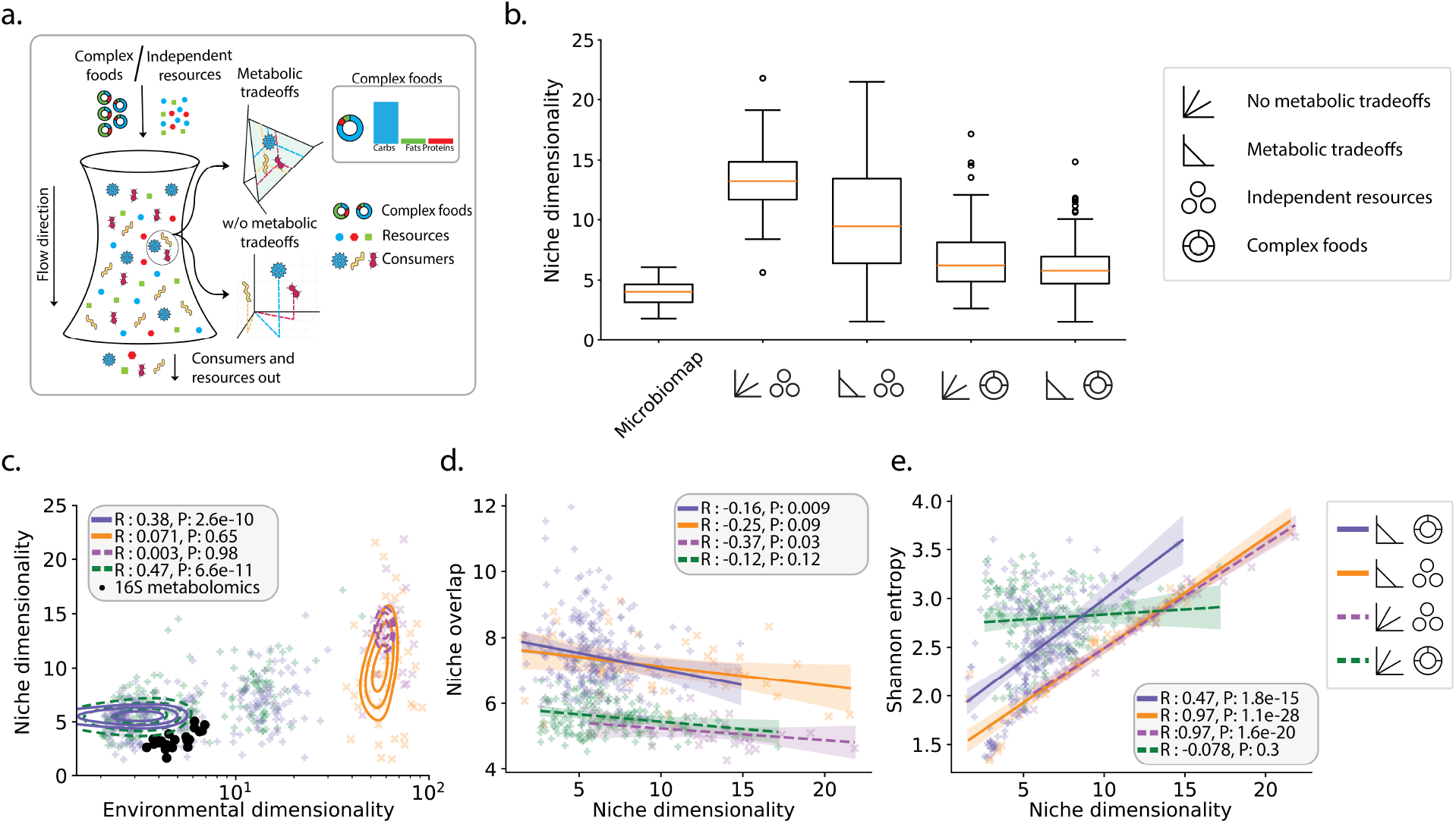
Tradeoffs generate low dimensionality and empirical covariation. **a**, CRM schematic and simulation strategy. **b**, Box plot of *η*_D_ across data and models: the Microbiomap collection^58^, shuffled data, baseline models reproducing diversity metrics, models with intrinsic metabolic tradeoffs, models with extrinsic environmental tradeoffs, and models with both tradeoffs, respectively (brackets represent 25^th^–75^th^ percentiles). **c–e**, Only the combined-tradeoff model recapitulates observed correlations between *η*_D_ and (c) environmental complexity, (d) metabolic niche overlap, and (e) *α*-diversity.

### C. Tradeoffs explain empirical trends

To identify mechanisms producing low *η*_D_ and its covariation with competition and diversity (Fig. 2), we simulated consumer–resource models (CRMs)^42,45^ (Appendix V; Fig. 3a). Given the strong correlation between *α*-diversity and niche dimensionality (Fig. 2c), we first investigated whether reproducing community diversity metrics was sufficient to explain the observed variation in *η*_D_. To that end, we selected 50 random microbiome studies from the ∼ 200 datasets in the Microbiomap collection^58^. The diversity metrics of this subset were statistically indistinguishable from the bigger collection (see Appendix V E). We performed Markov chain Monte Carlo sampling in model hyperparameters (e.g., magnitude of the consumer preference matrix, outflow rates, etc.) to match empirical mean *α*-diversity (Shannon and richness) and *β*-diversity (average pairwise Jensen–Shannon divergence). While the simulated microbiomes reproduced diversity metrics with an average error of less than 10%, their median niche dimensionality was *η*_D_ = 13.2 [11.7, 14.9] (brackets show 25^th^ and 75^th^ percentiles), significantly higher than the range of niche dimensionalities observed in real microbiomes (median *η*_D_ = 4.0 [3.2, 4.6]) (Figure 3b).

The *niche dimensionality hypothesis* posits that tradeoffs enable high diversity in low dimensional niche spaces^25–27,59^. In microbiomes, tradeoffs may arise from intrinsic constraints, including genomic streamlining (bias toward gene loss)^65^ or physiological limitations (restricted capacity to express multiple metabolic pathways)^66^. Following previous work^9,11,12,67^, we modeled *intrinsic* metabolic tradeoffs by limiting the ability of species to excel at utilizing multiple resources towards growth (see Appendix V). In simulations tuned to reproduce empirical diversity metrics, intrinsic tradeoffs enabled species to coexist in lower-dimensional niches while preserving richness (Fig. 3b) (median *η*_D_ = 9.5 [6.4, 13.4]). However, even with metabolic tradeoffs, simulated niche dimensionalities remained significantly higher than those inferred from real microbiomes.

A second, often overlooked class of tradeoffs relevant to gut microbiomes are *extrinsic* tradeoffs imposed by host diet. Humans consume nutrients in the form of food groups (e.g., starches, meats, fruits) under roughly constant total calorie intake, so greater consumption of one group reduces intake of others^68,69^. This covariation generates correlations among metabolite concentrations that limit microbial access to independent nutrient axes, for example, constraining the simultaneous availability of carbohydrates and amino acids. Such environmentally induced tradeoffs effectively reduce the number of independent resource axes (Appendix VI), thereby lowering the niche dimensionality.

To investigate the effects of environmental tradeoffs, we simulated CRMs in which nutrients were supplied as correlated “food groups” (Appendix V; Fig. 3a). Analogous to the intrinsic metabolic tradeoffs, the relative abundances of individual food groups were constrained to a simplex, representing a constant total nutrient intake. Incorporating extrinsic environmental tradeoffs markedly reduced the inferred dimensionality (median *η*_D_ = 6.2 [4.9, 8.1]). Combining both intrinsic and extrinsic tradeoffs further lowered *η*_D_ (median *η*_D_ = 5.8 [4.7, 7.0]).

Crucially, only models with *both* tradeoffs reproduced the full empirical pattern set: *η*_D_ increased with environmental dimensionality, decreased with metabolic niche overlap, and increased with *α*-diversity (Fig. 3c–e). The positive coupling between environmental complexity and *η*_D_ emerged *only* when environmental tradeoffs were present.

### D. Host diet and health modulate microbial niche dimensionality

Because niche dimensionality reflects the complexity of the microbial environment, we hypothesized that host-level factors such as diet and health^3^ should systematically influence *η*_D_.

To directly test dietary effects on niche dimensionality (Appendix II), we analyzed longitudinal microbiome data from mice fed either a high-fat, high-sugar (HFHS) diet or a low-fat, plant-polysaccharide (LFPP) diet^70^. The HFHS diet provides a narrow set of rapidly digestible nutrients, whereas the LFPP diet supplies a diverse pool of fermentable polysaccharides that yield a chemically rich oligosaccharide environment. Consistent with this difference in nutrient complexity, LFPP-fed mice exhibited significantly higher *η*_D_ (Carmody^70^, Mann-Whitney U test p-value: 5.8 × 10^−35^) than HFHS-fed mice (Fig. 4a). Controlled human dietary studies show similar effects: animal-based diets reduce metabolomic complexity, whereas plant-rich diets expand it^70,71^. In agreement, cross-sectional analysis of human infant microbiomes^72^ (Fig. 4a) revealed that infants consuming complementary plant-based foods in addition to formula had higher *η*_D_ than infants fed exclusively on formula (He^72^, Mann-Whitney U test p-value: 7.3 × 10^−79^). Together, these results demonstrate that dietary complexity expands the dimensionality of microbial niche space.

**FIG. 4.**
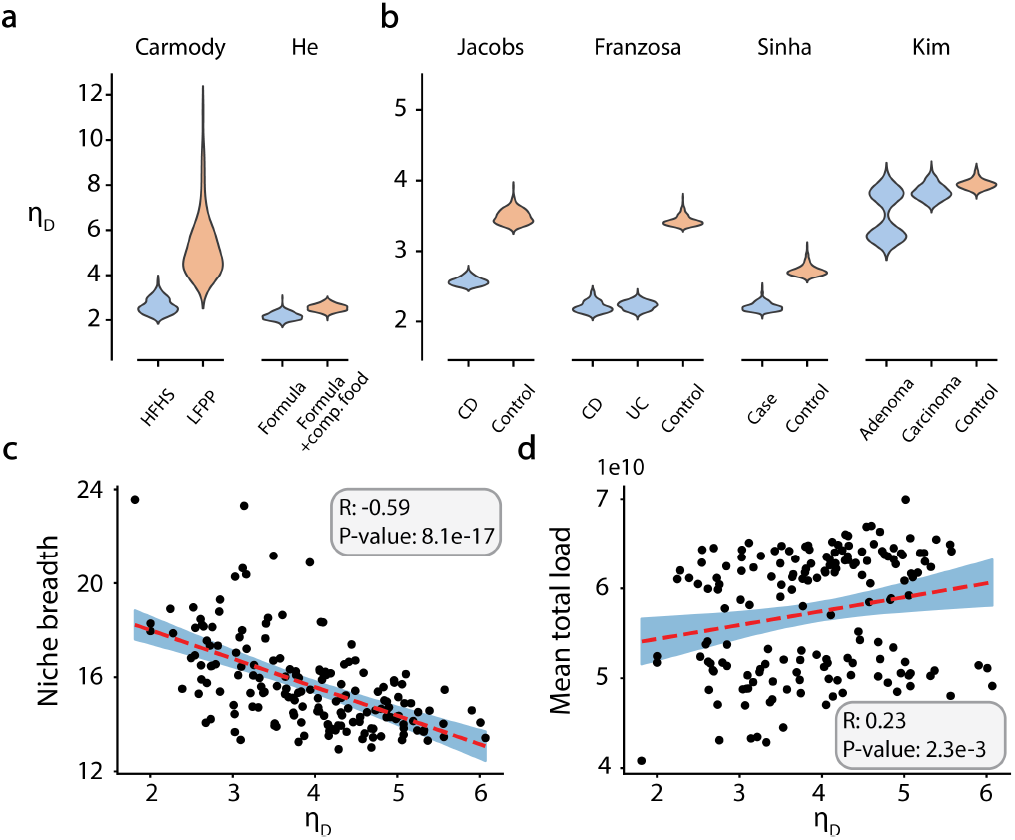
Niche dimensionality tracks diet complexity and health. **a**, Longitudinal mouse microbiomes: higher *η*_D_ under LFPP (diverse) vs. HFHS (narrow) diets; human infants: higher *η*_D_ with complementary foods vs. formulaonly. **b**, Higher *η*_D_ in healthy vs. inflammatory bowel disease or colorectal cancer. **c**, *η*_D_ correlates negatively with niche breadth (generalists’ prevalence). **d**, *η*_D_ correlates positively with predicted total microbial load.

Health status showed similar modulation. Across cross-sectional studies of inflammatory bowel disease^73,74^ and colorectal cancer^75,76^, healthy controls consistently exhibited higher *η*_D_ than patients (Fig. 4b) (Mann-Whitney U test P-values, Jacobs^73^: 1.1 × 10^−99^, Franzosa^74^: 1.1 × 10^−99^ in both cases, Sinha^75^: 1.1 × 10^−99^, Kim^76^: 1.2 × 10^−34^ for Control vs. Adenoma and 7.3 × 10^−78^ for Control vs Carcinoma). More broadly, dimensionality covaried with indicators of stress: it correlated negatively with niche breadth (mean number of resources consumbed by bacteria)^60^ (Spearman *r* = − 0.59, *p* = 2.4 × 10^−17^; Fig. 4c) and positively with total microbial load^61^, estimated using a recently developed machine learning model^77^ (Spearman *r* = 0.23, *p* = 2.3 × 10^−3^; Fig. 4d) (Appendices III, II). Together, these analyses identify reduced *η*_D_ as a quantitative hallmark of stressed or diet-simplified microbiomes and demonstrate that host-level factors systematically shape the effective ecological complexity experienced by gut communities.

## III. DISCUSSION

A central challenge in ecology is to explain the origins of biodiversity. Within niche theory, two contrasting coexistence regimes are often invoked^39^: ordered coexistence maintained in low-dimensional niche spaces^25,59^ and coexistence emerging from very high-dimensional interactions^78,79^. The former predicts a direct link between dimensionality, competition, and biodiversity—the *niche dimensionality hypothesis*^25–27^.

Across a large compendium of host-associated microbiomes, we find strong empirical evidence that niche dimensionality is a central organizing variable in microbial ecosystems. Reduced dimensionality was consistently associated with greater metabolic niche overlap, lower diversity, higher prevalence of generalists, and reduced microbial load. These relationships reveal that the ecological principles governing coexistence in macroecology extend naturally to microbial ecosystems.

Simulations using the consumer/resource models reproduced these empirical relationships only when two types of tradeoffs were imposed. Species-intrinsic metabolic tradeoffs, arising from genetic or physiological limitations, restrict efficient use of multiple resources^9,11,67^. Environmentally imposed tradeoffs, in contrast, emerge from correlations among resource inflows under the constraint of constant nutrient intake, as in host diets that deliver metabolites in co-occurring food groups. The latter exerted the dominant effect: correlated resource supplies restricted the number of independent ecological axes, lowering *η*_D_. Although these tradeoffs were introduced solely to reproduce empirical variation in dimensionality, the resulting models naturally predicted the observed dependencies among *η*_D_, environmental complexity, competition, and diversity. This concordance supports the interpretation that tradeoffs, particularly environmentally imposed correlations, govern the low-dimensional organization of microbial ecosystems.

Tradeoffs substantially reduced the median *in silico* niche dimensionality from 13.2 to 5.8, yet a small residual difference remained compared to empirical microbiomes (median *η*_D_ = 4). Other mechanisms that correlate environmental variables, such as niche creation through crossfeeding^14–16,80^ or coupled abiotic factors (pH, temperature, etc.)^81^, may also lower dimensionality. Exploring these effects is an important direction for future work.

These findings highlight a limitation of many existing microbiome models^42,45–49^, which reproduce community-level statistics but implicitly assume extremely high-dimensional niches (for example, by assuming independently sampled interactions between species^39^), effectively removing constraints imposed by niche dimensionality. Consistent with this, randomly initialized consumer–resource models captured species abundance distributions but failed to reproduce the low *η*_D_ observed empirically. Our work shows that future theoretical work must explicitly incorporate dimensionality constraints. In consumer–resource formulations, low dimensionality arises from correlated resource supplies; in generalized Lotka–Volterra systems, it may manifest as a low-rank interaction matrix^20–22,39^. Notably, empirical analyses in plants^38^ and microbes^82^ likewise reveal structured competition matrices, suggesting that dimensional constraints on coexistence represent a general ecological principle.

Efforts to coarse-grain microbial community composition, whether by linking it to aggregate functions^83–85^ or by capturing its fluctuations through low-dimensional modes^13,86^, reflect a growing recognition that ecological complexity may often collapse onto a few dominant axes. Comparable dimensional compression has been observed across biological scales, from correlated neural activity to protein mutational landscapes^87,88^. Notably, our own work revealed that the consumer–resource model can be recast as a latent-variable system^55,56^, suggesting that low dimensionality may represent a unifying principle governing the organization of diverse biological systems^89^.

In summary, microbial community structure can be understood through the lens of niche dimensionality. Both empirical data and mechanistic simulations demonstrate that dimensional constraints, mediated by metabolic and environmental tradeoffs, govern coexistence in microbiomes. By quantifying these constraints directly from abundance data, our framework links environmental complexity, biodiversity, and ecosystem health, establishing a quantitative bridge between microbial ecology and the long-standing traditions of niche theory in macroecology.

## Appendix

All scripts used in this paper can be found at https://github.com/karthik-yale/niche-dimensionality.

### I. APPENDIX: JOINT SPECIES DISTRIBUTION MODEL

Traditional joint species distribution models (JSDMs) model species abundances as functions of measured environmental covariates^50–53^, capturing the portion of ecological variation explained by known/measured factors. In contrast, their latent-variable extensions^53–56^ invoke unobserved environmental axes that capture shared, unmeasured, or unknown environmental influences shaping community composition.

Here, we use a latent variable JSDM that can be derived from the consumer/resource model^54–56^. Specifically, the relative abundance of taxons are modeled as a non-linear low-rank matrix factorization:

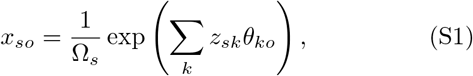

where *x*_*so*_ is the relative abundance of taxon *o* in sample *s, z*_*sk*_ are sample-specific latent embeddings (latent “resources”), and *θ*_*ko*_ are species-specific latent embeddings (latent preferences).

#### A. Fitting the model to data

We consider that we have measured abundances *n*_*so*_ of species *o* ∈ [1, *O*] in sample *s* ∈ [1, *S*]. We assume that the total read count *N* = ∑ _*o*_ *n*_*so*_ is constant across all samples. The multinomial likelihood of the data ***n*** given the model parameters ***z*** and ***θ*** is given by

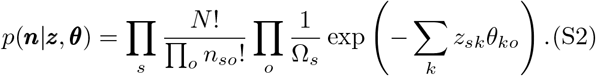

The log-likelihood ℒ (up to an additive and a multiplicative constant) is given by

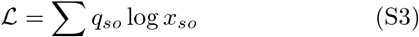

where *q*_*so*_ = *n*_*so*_*/N* are the observed relative abundances. We obtain the latent embeddings using a maximum likelihood approach. The gradients are given by:

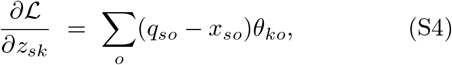

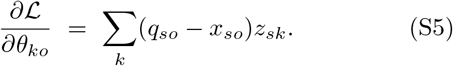

We fit the model to the data using simple gradient descent. The learning procedure is stopped when the sum of the magnitudes of the relative gradients falls below a threshold, 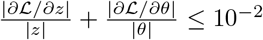 .

#### B. Estimating niche dimensionality

Niche dimensionality was calculated based on the rate of decrease of the KL-divergence between data and fit (error function for the gradient descent) with increasing dimension of the latent space (*K*). To capture the rate of decrease, KL-divergence for *K* = 1 to 5 was fit with an exponential function:

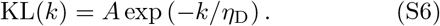

The coefficient *η*_D_ in the exponent was defined to be the niche dimensionality. The fit to the exponential function was performed by log-transforming the KL-divergences and using linear regression based on L2-norm.

### II. APPENDIX: DATA CURATION

#### A. The Microbiomap collection

Microbiomap is a large collection of human microbiome datasets, processed uniformly across ∼ 400 different datasets comprising ∼ 1.7 × 10^5^ independent samples^58^. Individual datasets had varying samples (ranging from ∼ 10 to ∼ 1000), very different number of bacterial ASVs, and total read count. To reduce potential biases related to sample size, number of bacteria, and read counts, we randomly sub-sampled all datasets to 5000 reads 10 times to create 10 variants. To eliminate the bias arising from varying number of samples, we subsampled all datasets to 100 samples (we only included datasets with more than 100 samples). This process was also repeated 10 times. Therefore, there were 100 total replicates for each project. To minimize technical noise in abundance measurements^90^, we only included bacteria whose average relative abundance across samples was *>* 0.1%. After this filtering, only the datasets with > 70 ASVs were used for further analysis.

The total load for each sample (Figure 4d) was estimated using a recent machine learning method^77^. The total load for a dataset was reported as the average across samples.

#### B. Paired 16S/metabolomics data

We used a large curated collection of paired 16S microbiome composition and metabolomics datasets^57^ to investigate the relationship between metabolomics and niche dimensionality. Metabolomics data were log-transformed and z-scored to standardize different metabolites. Metabolomics datasets often have many zeros. To avoid taking logarithms of zeros, in every dataset, we offset all metabolite measurements by the tenth percentile of the non-zero abundance prior to logtransformation.

Niche dimensionality for the microbiome was computed as described in the main text. The dimensionality of the metabolic environment was computed by performing singular value decomposition on the log-transformed metabolomics data and fitting an exponential decay to the singular value spectrum. To avoid artifacts related to sample size, we sub-sampled each dataset to 50 samples (or 70% of the sample size if the sample size was lower than 50). Each dataset was sub-sampled randomly to the value determined by the above procedure 100 times. The quantities of interest (niche dimensionality and environmental complexity) for each dataset was then calculated as the mean of that across the 100 replicates.

##### 1 Predicting metabolomics using learned ecosystem embeddings

Each dataset was sub-sampled 20 times wherein each time 50 random samples (or 70% of the total number of samples if the total number of samples was less than 50) were selected. This dataset was fit with the latent variable JSDM with *K* = 10 latent variables. The following procedure was performed on this learned set of latent variables and a shuffled set where the shuffling was performed for each latent variable separately. This preserved the distribution of each latent variable across samples. The latent variables were then split into training and testing sets with a ratio of 80% and 20% respectively. A linear model was trained on the training set to fit to the corresponding metabolite concentrations. The performance of latent variable predicting the metabolite concentrations was quantified as the difference in out of sample Pearson *R*^2^ of the top 10% of best predicted metabolites. The Pearson *R*^2^ for the same metabolites predicted by the shuffled latent variables with their own linear model served as the null model.

#### C. Host health and diet

The datasets studying host health and diet were processed as follows. To eliminate sample size effect, each group in a study was randomly sub-sampled to 20 samples. Niche dimensionality was calculated on this smaller dataset of 20 samples and the process was repeated 300 times. The infant milk study^72^ and the murine micorbiome diet study^70^ were a mix of longitudinal and crosssectional data. This was handled in infant milk study by grouping samples from a subject and replacing it with its mean, effectively making the dataset cross-sectional. In the murine diet study, niche dimensionality was determined for the longitudinal microbiome data for each mouse separately.

### III. APPENDIX: EVALUATING NICHE OVERLAP USING FLUX BALANCE ANALYSIS

#### A. Obtaining the metabolite preference matrix

Microbial species survival in the gut largely depends on the ability of species to metabolize available nutrients for growth^42,91^. To approximate the metabolic niches of gut bacteria, we analyzed thousands of manually and computationally curated genome-scale metabolic models^62^ using flux balance analysis (FBA)^63^. Although FBA does not yield quantitative preference parameters, it reliably predicts whether a species can utilize a particular resource^64^.

To that end, we identified *resources* as annotated exchange reactions. In any particular model (representing a bacterial species, and oftentimes a specific bacterial strain), we first allowed all exchange reactions to have an influx of a fixed amount and maximized the growth rate (biomass production). We then individually increased the upper limit of influx for each exchange reaction and calculated maximum growth rate. The nutrients that were able to increase the growth rate were deemed to be those that the model could utilize for growth. All calculations were performed anaerobically (setting influx of molecular oxygen to zero). We did not differentiate between resources that contributed carbon, nitrogen, sulfur, etc. At the end of the calculation, we obtained a binary matrix that represented predicted abilities (presence/absence) of individual models to utilize individual metabolite resources.

#### B. Mapping flux balance analysis models to taxa

For every sub-sampled dataset (see above), we aimed to match the ASVs to the ∼ 7000 genome scale metabolic models. To do so, we used taxonomy in a hierarchical manner. For every ASV, we matched the closed finegrained taxonomic label to that of the genome scale models. We only allow mapping to the class level. Any mapping at a broader taxonomic resolution was discarded. This way, each ASV was potentially mapped to multiple metabolic models (corresponding to multiple consumer/preference vectors). For calculating the niche overlap and mean number of resources consumed, we randomly chose one of these vectors.

#### C. Niche overlap calculation

Niche overlap in the consumer-resource model was calculated as follows. Each organism (consumer) was represented as a binary vector of resource preferences. For an average *k* − tuple of organisms *k* ∈ [2, 6], we first estimated the number of resources consumed by *all k* organisms by first randomly drawing *k* organisms from the ecosystem and then averaging over multiple draws. The upper limit of *k* = 6 was chosen due to computational constraints arising from combinatorial nature of this calculation.

Due to the sparsity of the resource preference matrix, the average overlap between *k* species decreases with increasing *k*. The null expectation for this decrease for a given sparsity *ρ* of the consumer preference matrix and the number of resources *M* is simply *Mρ*^*k*^. For each *k*, we computed the competition as the ratio of the estimated overlap to the null expectation. The total niche overlap was defined as the sum over all values of *k*.

The same procedure was followed for niche overlap calculation in microbiomap with only the null model being calculated differently. The null model for microbiomap was simply the niche overlap calculation performed from a random set of bacteria selected to have a similar consumer preference matrix sparsity.

##### 1. Niche breadth

In any given dataset, the niche breadth (Figure 4c) was estimated as the mean number of resources consumed by bacteria (after mapping bacteria to resources, see above). As above, only bacteria with mean abundance > 0.1% were included in the analysis.

##### 2. Predicting species-preferences using learned taxon loadings

For each replicate of a dataset, the latent variable JSDM was trained with 20 latent variables. The species loadings *θ*_*ko*_ learned from this procedure was used as a regressor in a logistic regression model to predict the average consumer preference matrix constructed using the mapping to FBA models described above. The plot is a histogram of the difference in the log-likelihood of the model fit of all resources across all taxa used the learnt *θ* and corresponding shuffled *θ*.

### IV. APPENDIX: ALTERNATIVE METRICS TO COMPUTE NICHE DIMENSIONALITY

To further validate our approach to compute niche dimensionality (Figure 1), we compared estimated *η*_D_ to other standard metrics to infer matrix dimensionality. Specifically, we chose participation ratio^92^ and spectrum entropy^93^. The first step in determining these two metrics for any matrix is computing the singular value decomposition to obtain singular values {*λ*_*i*_ }. Participation ratio (PR) and spectrum entropy (SE) are then defined as

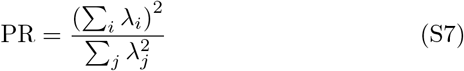

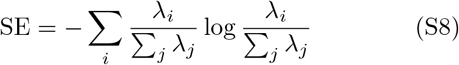

**FIG. S1.**
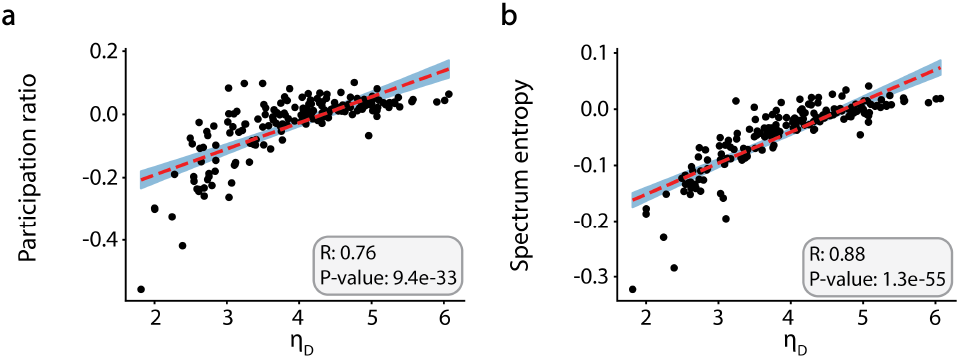
Participation ratio and spectrum entropy correlate strongly with estimated *η*_D_.

We compute the two metrics by first performing latent variable inference with a large latent dimension (*K* = 20) and then performing singular value decomposition on the matrix *M* = *Z* × Θ.

Both these metrics were corrected by a null model. Corrected metrics correlated very strongly with our estimate of the niche dimensionality (with a Spearman R of 0.76 and 0.88 respectively, Figure S1). Null model correction was done as follows. The metric of choice is calculated for the regular relative abundance data and for the relative abundance data shuffled such that the relative abundance of each taxon is shuffled across samples after which each sample is re-normalized to sum up to 1. The null model corrected metric is then the difference between the two.

### V. APPENDIX: CONSUMER-RESOURCE MODEL SIMULATIONS

#### A. Solving the consumer-resource model (CRM) as a convex optimization problem

The consumer resource model simulations were performed as convex optimization problems similar to the community simulator package by Marsland et al.^94^.

We first start with the equations describing the CRM:

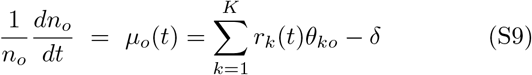

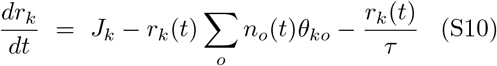

The convex optimization problem to determine community composition is stated as follows. We want to minimize the Kullback-Leibler divergence between *J*_*k*_*/δ* and *r*_*k*_ given by,

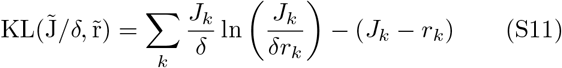

subject to the constraint that *µ*_*o*_(*t*) ≤ 0. The consumer populations are the Lagrange multipliers that impose the constraints.

#### B. Parameter selection

To explore the parameter space of the consumer resource models, we first look at the set of equations satisfied at steady state is given by,

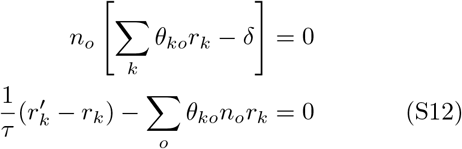

where we have rewritten the inflow rate in terms of the dilution rate 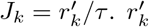 is the steady state value the resources would take in the absence of consumers. We define: 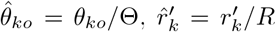 and 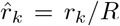 which are dimensionless quantities.

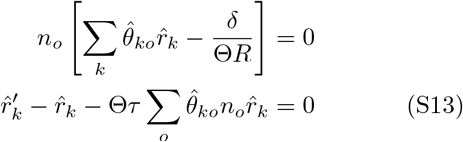

We see that just two combinations of parameters, *δ/*(Θ*R*) and Θ*τ*, capture the entirety of the variation at steady state once 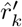 and 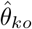 are fixed. 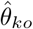 is chosen to be a uniform sparse matrix with a density of 10 percent and 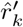 is sampled from a uniform distribution. To simulate the variation between samples within a dataset, we generate each sample’s inflow, 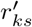 where *s* denotes the sample index, from a gaussian distribution centered at 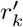 with a variance of 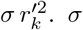, Θ and *δ/*Θ along with *ρ*, which will be introduced in the subsection V D, are parameters we optimize with simulated annealing.

#### C. Implementing intrinsic metabolic tradeoffs

For the simulations with intrinsic metabolic tradeoffs^9,11^, we normalized the randomly sampled consumer preference matrix, 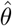, by the sum of each species’ preference vector, 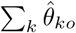 . As a result, the preference vectors for all species summed up to 1. This matrix was then scaled by Θ chosen by the simulated annealing procedure.

#### D. Implementing environmental tradeoffs

We capture the systematic variation of environmental influence on samples by grouping the presence/absence of resources. As an example, a typical person’s diet comprises multiple food groups (starches, proteins, fiber, etc.). These food groups have their own set of metabolites where some of these metabolites are present in only one group and some are shared by both groups.

In our simulations we enforce this structure by generating scaling vectors that alters the magnitude of the resource inflow rates across samples. These scaling vectors are generated as random linear combinations of different masking vectors 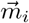, which are binary vectors that denote the presence/absence of a resource in a particular food group *i*. The scaling vector of sample *s*, denoted by 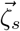, is given by,

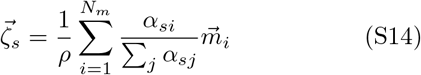

where *N*_*m*_ is the total number of masking vectors, *ρ* is the density of the masking vectors, 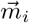 is the *i*^th^ masking vector, *α*_*si*_ are the relative abundances of individual food groups ( ∑_*i*_ *α*_*si*_ = 1), which are sampled from a uniform distribution for each sample and then normalized.

#### E. Simulation details

To investigate the role of metabolic and extrinsic tradeoffs, we randomly selected 50 datasets from the Microbiomap collection^58^. The niche dimensionality, *α* − diversity, species richness, and *β* − diversity of this subset was statistically indistinguishable compared to the larger set, with p-values of 0.26, 0.59, 0.51, and 0.26 respectively.

All the simulations in the main text started with 250 resources, 700 consumers, and a consumer preference matrix with 10% sparsity. The preferences values were uniformly distributed between 0 and Θ (a hyperparameter of the model). The mean resource inflow values were also sampled from a uniform distribution from 0 to *R* (another hyperparameter) which was set to *R* = 5. For the simulations with environmental tradeoffs (see above), the masking vectors together were sampled from a random sparse matrix with density *ρ* and the *α* matrix was populated with random samples from a uniform random distribution between 0 and 1.

Notably, the total number of resources did not affect our overall conclusions. Model simulations with 50 resources also reproduced low niche dimensionality distributions with tradeoffs and the observed empirical relationships between niche dimensionality and community structure. Simulations with 50 resources without metabolic tradeoffs can only support at best 50 consumers. Therefore, we only performed these calculations with metabolic tradeoffs.

**FIG. S2.**
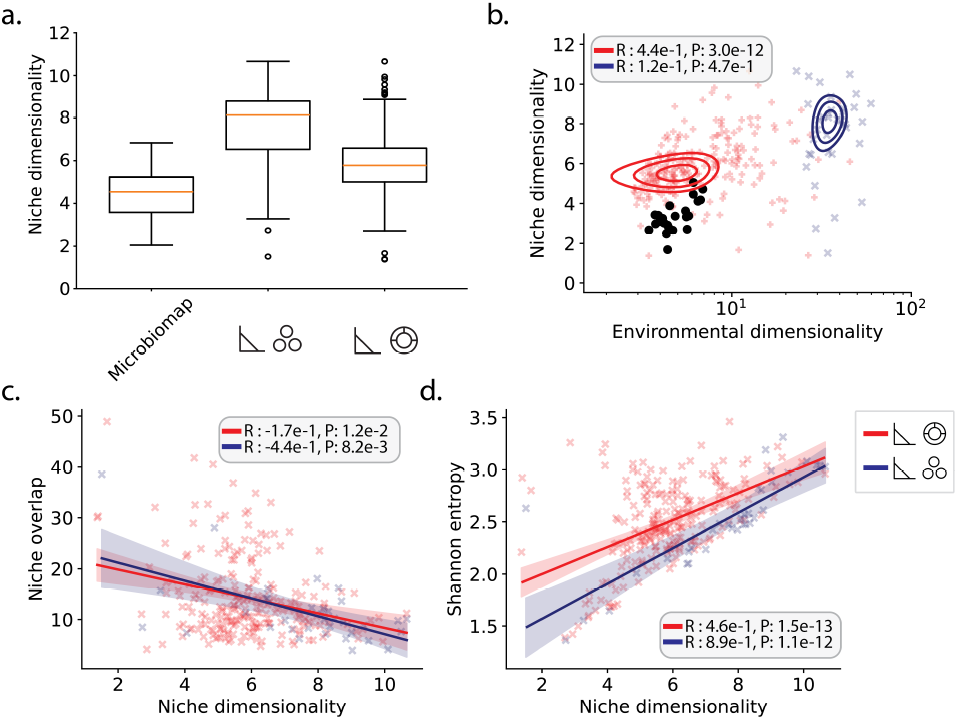
Niche dimensionality and correlations with consumer/resource simulations with 50 resources.

We pooled the results from 2-7 number of food groups. The distribution of niche dimensionality remained largely consistent above the seven food group threshold.

### VI. APPENDIX: RESOURCE COVARIATION AND NICHE DIMENSIONALITY

In this section we will show how environmental covariation reduces the dimensionality of niche overlaps. The calculation follows classic treatment by MacArthur^8,18,19^ that estimates niche overlap as Gaussian overlap integrals. Assume that there is a *D* dimensional resource space in which the *niche* of an organism is expressed a normal distribution,

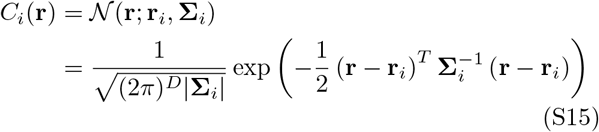

Here, *C*_*i*_(**r**) is the population size (or growth rate) of organism *i* if the niche variables take the value **r**.

Most standard niche overlap calculations assume that **r** are uniformly distributed^8,18,19^. Here, we explicitly account for correlations in the environment. To that end, let *S*(**r**) = 𝒩 (**r, r**_*R*_, **Σ**_*R*_) denote the distribution over environmental variables, assumed to be a multivariate Gaussian distribution with mean **r**_*R*_ and a covariance matrix **Σ**_*R*_.

We define the competition between two species *i* and *j* to be the overlap integral

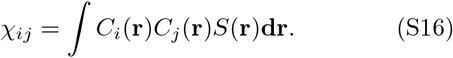

To evaluate the integral, we shift the coordinate frame so that the mean of the environmental variable distribution **r**_*R*_ is at the origin. We define means of the organism’s niche preference distribution form this frame to be **x**_*i*_ = **r**_*i*_ − **r**_*R*_ and **x**_*j*_ = **r**_*j*_ − **r**_*R*_, the pooled inverse covariance as 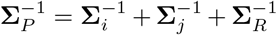 and the pooled mean 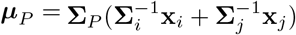.

The overlap integral can be expressed as:

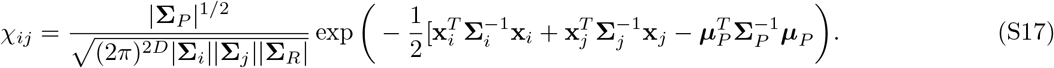

We now impose structure in the environment by assuming that the resources are sampled from a lower dimensional space (of dimension *k < D*). To do so, we assume **r** = **Bu** where **B** is a *D*× *k* matrix whose columns form an orthonormal basis. To work in the subspace we transform distributions in the *k*-dimensional space of **u**.

We have **u**_*R*_ = **0, Ω**_*R*_ = **B**^*T*^ **Σ**_*R*_**B, u**_*i*_ = **B**^*T*^ (**r**_*i*_ − **r**_*R*_) and **Ω**_*i*_ = **B**^*T*^ **Σ**_*i*_**B**. The transformed *k*-dimensional Gaussian PDFs are given by,

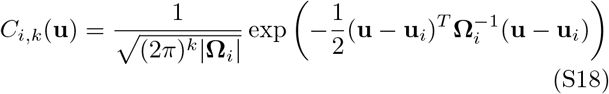

With these definitions, the niche overlap in the lowerdimensional space is given by

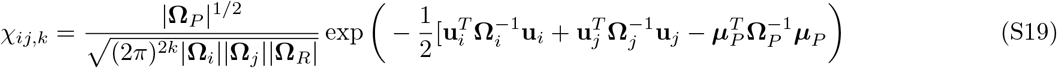

where 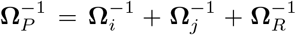 is the pooled inverse covariance and 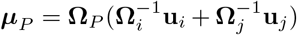 is the pooled mean.

**FIG. S3.**
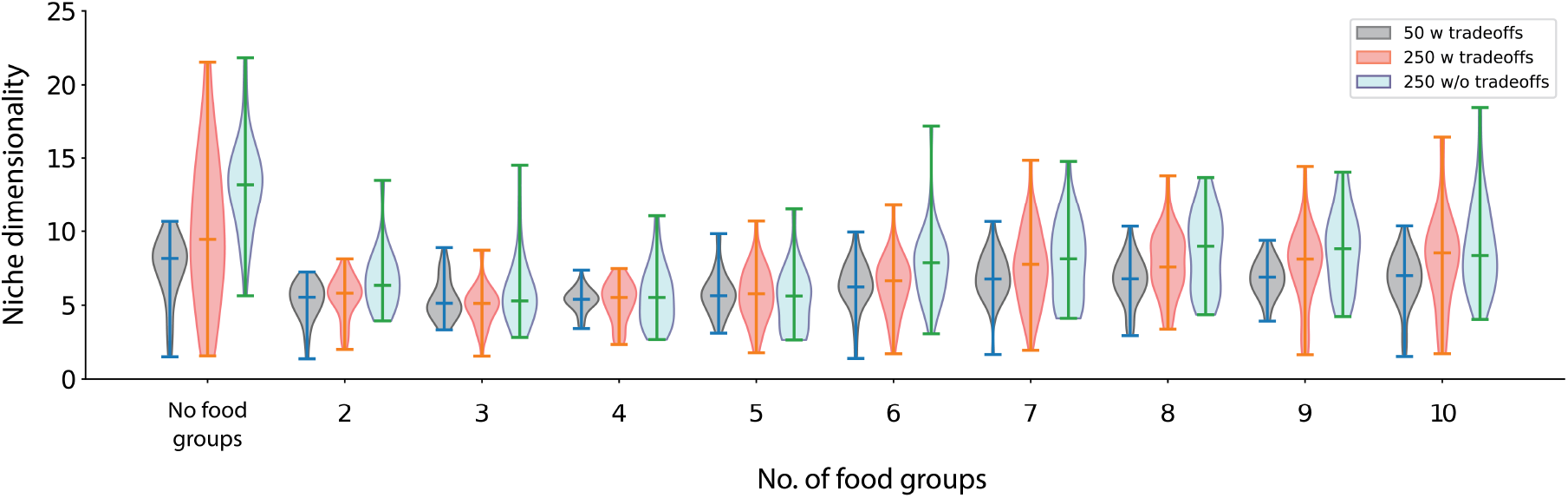
Violin plots of niche dimensionality versus number of food groups in the consumer/resource simulations.

A comparison of Eq. S17 and Eq. S19 shows that restricting the environments to a lower dimension leads to overlap matrices that are computed in a lower dimensional space as well^20,21,38,82^.

## References

1 P. J. Turnbaugh, R. E. Ley, M. Hamady, C. M. Fraser-Liggett, R. Knight, and J. I. Gordon, “The human microbiome project,” Nature 449, 804–810 (2007).

2 A. K. Nash, T. A. Auchtung, M. C. Wong, D. P. Smith, J. R. Gesell, M. C. Ross, C. J. Stewart, G. A. Metcalf, D. M. Muzny, R. A. Gibbs, et al., “The gut mycobiome of the human microbiome project healthy cohort,” Microbiome 5, 1–13 (2017).

3 J. A. Gilbert, M. J. Blaser, J. G. Caporaso, J. K. Jansson, S. V. Lynch, and R. Knight, “Current understanding of the human microbiome,” Nature medicine 24, 392–400 (2018).

4 I. Tapio, T. J. Snelling, F. Strozzi, and R. J. Wallace, “The ruminal microbiome associated with methane emissions from ruminant livestock,” Journal of animal science and biotechnology 8, 1–11 (2017).

5 J. M. Chase and M. A. Leibold, Ecological niches: linking classical and contemporary approaches (University of Chicago Press, 2009).

6 M. Leibold and V. F. Frans, “Re-revisiting the niche concept,” (2025).

7 G. E. Hutchinson, “Cold spring harbor symposia on quantitative biology,” (No Title) 22, 415 (1957).

8 R. MacArthur and R. Levins, “The limiting similarity, convergence, and divergence of coexisting species,” The american naturalist 101, 377–385 (1967).

9 A. Posfai, T. Taillefumier, and N. S. Wingreen, “Metabolic trade-offs promote diversity in a model ecosystem,” Physical review letters 118, 028103 (2017).

10 A. Goyal, V. Dubinkina, and S. Maslov, “Multiple stable states in microbial communities explained by the stable marriage problem,” The ISME journal 12, 2823–2834 (2018).

11 R. Caetano, Y. Ispolatov, and M. Doebeli, “Evolution of diversity in metabolic strategies,” Elife 10, e67764 (2021).

12 Z. R. Miller and J. P. O’Dwyer, “Metabolic trade-offs can reverse the resource-diversity relationship,” The American Naturalist 204, E85–E98 (2024).

13 A. V. Narla, T. Hwa, and A. Murugan, “Dynamic coexistence driven by physiological transitions in microbial communities,” Proceedings of the National Academy of Sciences 122, e2405527122 (2025).

14 M. Gralka, R. Szabo, R. Stocker, and O. X. Cordero, “Trophic interactions and the drivers of microbial community assembly,” Current Biology 30, R1176–R1188 (2020).

15 M. Dal Bello, H. Lee, A. Goyal, and J. Gore, “Resource–diversity relationships in bacterial communities reflect the network structure of microbial metabolism,” Nature Ecology & Evolution 5, 1424–1434 (2021).

16 S. Estrela, J. Diaz-Colunga, J. C. Vila, A. Sanchez-Gorostiaga, and A. Sanchez, “Diversity begets diversity under microbial niche construction,” BioRxiv, 2022–02 (2022).

17 E. R. Pianka, “Niche overlap and diffuse competition,” Proceedings of the National Academy of Sciences 71, 2141–2145 (1974).

18 R. H. MacArthur, Geographical ecology: patterns in the distribution of species (Princeton University Press, 1984).

19 R. MacArthur, “Species packing and competitive equilibrium for many species,” Theoretical population biology 1, 1–11 (1970).

20 R. M. Yoshiyama and J. Roughgarden, “Species packing in two dimensions,” The American Naturalist 111, 107–121 (1977).

21 C. Rappoldt and P. Hogeweg, “Niche packing and number of species,” The American Naturalist 116, 480–492 (1980).

22 T. J. Case, “Niche packing and coevolution in competition communities,” Proceedings of the National Academy of Sciences 78, 5021–5025 (1981).

23 E. R. Pianka, “The structure of lizard communities,” Annual review of ecology and systematics, 53–74 (1973).

24 R. Costa-Pereira, M. S. Araújo, F. L. Souza, and T. Ingram, “Competition and resource breadth shape niche variation and overlap in multiple trophic dimensions,” Proceedings of the Royal Society B 286, 20190369 (2019).

25 W. S. Harpole and D. Tilman, “Grassland species loss resulting from reduced niche dimension,” Nature 446, 791–793 (2007).

26 W. S. Harpole, L. L. Sullivan, E. M. Lind, J. Firn, P. B. Adler, E. T. Borer, J. Chase, P. A. Fay, Y. Hautier, H. Hillebrand, et al., “Addition of multiple limiting resources reduces grassland diversity,” Nature 537, 93–96 (2016).

27 E. T. Borer, J. B. Grace, W. S. Harpole, A. S. MacDougall, and E. W. Seabloom, “A decade of insights into grassland ecosystem responses to global environmental change,” Nature Ecology & Evolution 1, 0118 (2017).

28 N. Eisenhauer, W. Schulz, S. Scheu, and A. Jousset, “Niche dimensionality links biodiversity and invasibility of microbial communities,” Functional Ecology 27, 282–288 (2013).

29 P. Nosil and C. P. Sandoval, “Ecological niche dimensionality and the evolutionary diversification of stick insects,” PLoS one 3, e1907 (2008).

30 M. J. Larcombe, G. J. Jordan, D. Bryant, and S. I. Higgins, “The dimensionality of niche space allows bounded and unbounded processes to jointly influence diversification,” Nature Communications 9, 4258 (2018).

31 L.-M. Chevin, G. Decorzent, and T. Lenormand, “Niche dimensionality and the genetics of ecological speciation,” Evolution 68, 1244–1256 (2014).

32 J. E. Cohen, “Food webs and the dimensionality of trophic niche space,” Proceedings of the National Academy of Sciences 74, 4533–4536 (1977).

33 H. J. Morowitz, “The dimensionality of niche space,” Journal of Theoretical Biology 86, 259–263 (1980).

34 A. G. Rossberg, Food webs and biodiversity: foundations, models, data (John Wiley & Sons, 2013).

35 R. J. Williams and D. W. Purves, “The probabilistic niche model reveals substantial variation in the niche structure of empirical food webs,” Ecology 92, 1849–1857 (2011).

36 A. Eklöf, U. Jacob, J. Kopp, J. Bosch, R. Castro-Urgal, N. P. Chacoff, B. Dalsgaard, C. de Sassi, M. Galetti, P. R. Guimarães, et al., “The dimensionality of ecological networks,” Ecology letters 16, 577–583 (2013).

37 D. C. Laughlin, “The intrinsic dimensionality of plant traits and its relevance to community assembly,” Journal of ecology 102, 186–193 (2014).

38 D. B. Stouffer, O. Godoy, G. V. Dalla Riva, and M. M. Mayfield, “The dimensionality of plant–plant competition,” bioRxiv, 2021–11 (2021).

39 M. Barbier, C. De Mazancourt, M. Loreau, and G. Bunin, “Fingerprints of high-dimensional coexistence in complex ecosystems,” Physical Review X 11, 011009 (2021).

40 K. Z. Coyte, J. Schluter, and K. R. Foster, “The ecology of the microbiome: networks, competition, and stability,” Science 350, 663–666 (2015).

41 M. Tikhonov and R. Monasson, “Collective phase in resource competition in a highly diverse ecosystem,” Physical review letters 118, 048103 (2017).

42 R. Marsland III, W. Cui, and P. Mehta, “A minimal model for microbial biodiversity can reproduce experimentally observed ecological patterns,” Scientific reports 10, 3308 (2020).

43 W. Cui, R. Marsland III, and P. Mehta, “Diverse communities behave like typical random ecosystems,” Physical Review E 104, 034416 (2021).

44 J. Hu, D. R. Amor, M. Barbier, G. Bunin, and J. Gore, “Emergent phases of ecological diversity and dynamics mapped in microcosms,” Science 378, 85–89 (2022).

45 P.-Y. Ho, B. H. Good, and K. C. Huang, “Competition for fluctuating resources reproduces statistics of species abundance over time across wide-ranging microbiotas,” Elife 11, e75168 (2022).

46 J. Pasqualini, A. Maritan, A. Rinaldo, S. Facchin, E. Savarino, A. Altieri, and S. Suweis, “Microbiomes through the looking glass,” eLife 14 (2025).

47 X.-W. Wang and Y.-Y. Liu, “Origins of scaling laws in microbial dynamics,” Physical Review Research 5, 013004 (2023).

48 J. Camacho-Mateu, A. Lampo, M. Sireci, M. A. Muñoz, and J. A. Cuesta, “Sparse species interactions reproduce abundance correlation patterns in microbial communities,” Proceedings of the National Academy of Sciences 121, e2309575121 (2024).

49 R. Maskawa, H. Takayasu, L. Takayasu, W. Suda, and M. Takayasu, “Stochastic spatiotemporal growth model reproducing the universal statistical laws of the gut microbiome,” Physical Review Research 7, 013269 (2025).

50 O. Ovaskainen and N. Abrego, Joint species distribution modelling: With applications in R (Cambridge University Press, 2020).

51 L. J. Pollock, R. Tingley, W. K. Morris, N. Golding, R. B. O’Hara, K. M. Parris, P. A. Vesk, and M. A. McCarthy, “Understanding co-occurrence by modelling species simultaneously with a joint species distribution model (jsdm),” Methods in Ecology and Evolution 5, 397–406 (2014).

52 F. K. Hui, S. Taskinen, S. Pledger, S. D. Foster, and D. I. Warton, “Model-based approaches to unconstrained ordination,” Methods in Ecology and Evolution 6, 399–411 (2015).

53 D. I. Warton, F. G. Blanchet, R. B. O’Hara, O. Ovaskainen, S. Taskinen, S. C. Walker, and F. K. Hui, “So many variables: joint modeling in community ecology,” Trends in ecology & evolution 30, 766–779 (2015).

54 C. K. Fisher, T. Mora, and A. M. Walczak, “Variable habitat conditions drive species covariation in the human microbiota,” PLoS computational biology 13, e1005435 (2017).

55 M. Shahin, B. Ji, and P. D. Dixit, “Embed: Essential microbiome dynamics, a dimensionality reduction approach for longitudinal microbiome studies,” npj Systems Biology and Applications 9, 26 (2023).

56 G. Plata, K. Srinivasan, M. Krishnamurthy, L. Herron, and P. Dixit, “Designing host-associated microbiomes using the consumer/resource model,” mSystems 10, e01068–24 (2025).

57 E. Muller, Y. M. Algavi, and E. Borenstein, “The gut microbiome-metabolome dataset collection: a curated resource for integrative meta-analysis,” npj Biofilms and Microbiomes 8, 79 (2022).

58 R. J. Abdill, S. P. Graham, V. Rubinetti, M. Ahmadian, P. Hicks, A. Chetty, D. McDonald, P. Ferretti, E. Gibbons, M. Rossi, et al., “Integration of 168,000 samples reveals global patterns of the human gut microbiome,” Cell 188, 1100–1118 (2025).

59 D. Tilman, Resource competition and community structure, 17 (Princeton university press, 1982).

60 I. Veseli, Y. T. Chen, M. S. Schechter, C. Vanni, E. C. Fogarty, A. R. Watson, B. Jabri, R. Blekhman, A. D. Willis, M. K. Yu, et al., “Microbes with higher metabolic independence are enriched in human gut microbiomes under stress,” Elife 12, RP89862 (2025).

61 D. Vandeputte, G. Kathagen, K. D’hoe, S. Vieira-Silva, M. Valles-Colomer, J. Sabino, J. Wang, R. Y. Tito, L. De Commer, Y. Darzi, et al., “Quantitative microbiome profiling links gut community variation to microbial load,” Nature 551, 507–511 (2017).

62 A. Heinken, J. Hertel, G. Acharya, D. A. Ravcheev, M. Nyga, O. E. Okpala, M. Hogan, S. Magnuśdóttir, F. Martinelli, B. Nap, et al., “Genome-scale metabolic reconstruction of 7,302 human microorganisms for personalized medicine,” Nature Biotechnology 41, 1320–1331 (2023).

63 J. D. Orth, I. Thiele, and B. Ø. Palsson, “What is flux balance analysis?” Nature biotechnology 28, 245–248 (2010).

64 G. Plata, C. S. Henry, and D. Vitkup, “Long-term phenotypic evolution of bacteria,” Nature 517, 369–372 (2015).

65 A. Mira, H. Ochman, and N. A. Moran, “Deletional bias and the evolution of bacterial genomes,” TRENDS in Genetics 17, 589–596 (2001).

66 M. Basan, S. Hui, H. Okano, Z. Zhang, Y. Shen, J. R. Williamson, and T. Hwa, “Overflow metabolism in escherichia coli results from efficient proteome allocation,” Nature 528, 99–104 (2015).

67 E. Blumenthal and P. Mehta, “Geometry of ecological coexistence and niche differentiation,” Physical Review E 108, 044409 (2023).

68 K. L. Keller, J. Kirzner, A. Pietrobelli, M.-P. St-Onge, and M. S. Faith, “Increased sweetened beverage intake is associated with reduced milk and calcium intake in 3-to 7-year-old children at multi-item laboratory lunches,” Journal of the American Dietetic Association 109, 497–501 (2009).

69 A. B. Bontrager Yoder and D. A. Schoeller, “Fruits and vegetables displace, but do not decrease, total energy in school lunches,” Childhood Obesity 10, 357–364 (2014).

70 R. N. Carmody, G. K. Gerber, J. M. Luevano, D. M. Gatti, L. Somes, K. L. Svenson, and P. J. Turnbaugh, “Diet dominates host genotype in shaping the murine gut microbiota,” Cell host & microbe 17, 72–84 (2015).

71 L. A. David, C. F. Maurice, R. N. Carmody, D. B. Gootenberg, J. E. Button, B. E. Wolfe, A. V. Ling, A. S. Devlin, Y. Varma, M. A. Fischbach, et al., “Diet rapidly and reproducibly alters the human gut microbiome,” Nature 505, 559–563 (2014).

72 X. He, M. Parenti, T. Grip, B. Lönnerdal, N. Timby, M. Domellöf, O. Hernell, and C. M. Slupsky, “Fecal microbiome and metabolome of infants fed bovine mfgm supplemented formula or standard formula with breast-fed infants as reference: a randomized controlled trial,” Scientific reports 9, 11589 (2019).

73 J. P. Jacobs, M. Goudarzi, N. Singh, M. Tong, I. H. McHardy, P. Ruegger, M. Asadourian, B.-H. Moon, A. Ayson, J. Borneman, et al., “A disease-associated microbial and metabolomics state in relatives of pediatric inflammatory bowel disease patients,” Cellular and molecular gastroenterology and hepatology 2, 750–766 (2016).

74 E. A. Franzosa, A. Sirota-Madi, J. Avila-Pacheco, N. Fornelos, H. J. Haiser, S. Reinker, T. Vatanen, A. B. Hall, H. Mallick, L. J. McIver, et al., “Gut microbiome structure and metabolic activity in inflammatory bowel disease,” Nature microbiology 4, 293–305 (2019).

75 R. Sinha, J. Ahn, J. N. Sampson, J. Shi, G. Yu, X. Xiong, R. B. Hayes, and J. J. Goedert, “Fecal microbiota, fecal metabolome, and colorectal cancer interrelations,” PloS one 11, e0152126 (2016).

76 M. Kim, E. Vogtmann, D. A. Ahlquist, M. E. Devens, J. B. Kisiel, W. R. Taylor, B. A. White, V. L. Hale, J. Sung, N. Chia, et al., “Fecal metabolomic signatures in colorectal adenoma patients are associated with gut microbiota and early events of colorectal cancer pathogenesis,” MBio 11, 10–1128 (2020).

77 S. Nishijima, E. Stankevic, O. Aasmets, T. S. Schmidt, N. Nagata, M. I. Keller, P. Ferretti, H. B. Juel, A. Fullam, S. M. Robbani, et al., “Fecal microbial load is a major determinant of gut microbiome variation and a confounder for disease associations,” Cell 188, 222–236 (2025).

78 N. J. Kraft, O. Godoy, and J. M. Levine, “Plant functional traits and the multidimensional nature of species coexistence,” Proceedings of the National Academy of Sciences 112, 797–802 (2015).

79 J. S. Clark, D. Bell, C. Chu, B. Courbaud, M. Dietze, M. Hersh, J. HilleRisLambers, I. Ibášez, S. LaDeau, S. McMahon, et al., “High-dimensional coexistence based on individual variation: a synthesis of evidence,” Ecological Monographs 80, 569–608 (2010).

80 J. E. Goldford, N. Lu, D. Bajić, S. Estrela, M. Tikhonov, A. Sanchez-Gorostiaga, D. Segre, P. Mehta, and A. Sanchez, “Emergent simplicity in microbial community assembly,” Science 361, 469–474 (2018).

81 M. Sireci, M. A. Muñoz, and J. Grilli, “Environmental fluctuations explain the universal decay of species-abundance correlations with phylogenetic distance,” Proceedings of the National Academy of Sciences 120, e2217144120 (2023).

82 S. P. Mizrahi, H. Lee, A. Goyal, E. Owen, and J. Gore, “Structured interactions explain the absence of keystone species in synthetic microcosms,” bioRxiv, 2025–03 (2025).

83 J. Bergelson, M. Kreitman, D. A. Petrov, A. Sanchez, and M. Tikhonov, “Functional biology in its natural context: A search for emergent simplicity,” Elife 10, e67646 (2021).

84 J. Moran and M. Tikhonov, “Emergent predictability in microbial ecosystems,” arXiv preprint arXiv:2403.19372 (2024).

85 K. K. Lee, S. Liu, K. Crocker, J. Wang, D. R. Huggins, M. Tikhonov, M. Mani, and S. Kuehn, “Functional regimes define soil microbiome response to environmental change,” Nature, 1–11 (2025).

86 D. R. Hekstra and S. Leibler, “Contingency and statistical laws in replicate microbial closed ecosystems,” Cell 149, 1164–1173 (2012).

87 M. C. Morrell, A. J. Sederberg, and I. Nemenman, “Latent dynamical variables produce signatures of spatiotemporal criticality in large biological systems,” Physical review letters 126, 118302 (2021).

88 C. J. Russo, K. Husain, and A. Murugan, “Soft modes as a predictive framework for low-dimensional biological systems across scales,” Annual Review of Biophysics 54 (2025).

89 V. Thibeault, A. Allard, and P. Desrosiers, “The low-rank hypothesis of complex systems,” Nature Physics 20, 294–302 (2024).

90 B. W. Ji, R. U. Sheth, P. D. Dixit, Y. Huang, A. Kaufman, H. H. Wang, and D. Vitkup, “Quantifying spatiotemporal variability and noise in absolute microbiota abundances using replicate sampling,” Nature Methods 16, 731–736 (2019).

91 P.-Y. Ho, T. H. Nguyen, J. M. Sanchez, B. C. DeFelice, and K. C. Huang, “Resource competition predicts assembly of gut bacterial communities in vitro,” Nature Microbiology 9, 1036–1048 (2024).

92 P. Gao, E. Trautmann, B. Yu, G. Santhanam, S. Ryu, K. Shenoy, and S. Ganguli, “A theory of multineuronal dimensionality, dynamics and measurement,” BioRxiv, 214262 (2017).

93 O. Roy and M. Vetterli, “The effective rank: A measure of effective dimensionality,” in 2007 15th European signal processing conference (IEEE, 2007) pp. 606–610.

94 R. Marsland, W. Cui, J. Goldford, and P. Mehta, “The community simulator: A python package for microbial ecology,” Plos one 15, e0230430 (2020).

